# DNA-barcoded microneedle array patch enabling the profiling of in vivo spatiotemporal transcriptomics

**DOI:** 10.1101/2025.11.25.690619

**Authors:** Yuanbin Guo, Yile Zhang, Xinyin Li, Jingwen Kuang, Ke Cao, Feng Chen, Chunhai Fan, Yongxi Zhao

**Author notes:** Corresponding author: Yongxi Zhao, Feng Chen.

## Abstract

Recent years have witnessed profound advances in spatial transcriptomics (ST). However, current ST strategies are limited to tissue sections, unable to directly profile spatiotemporal transcriptome landscapes of individual living animals. To overcome this limitation, we present acupuncture-extracted in vivo spatiotemporal transcriptomics (aivST). It is mainly based on engineering clinical-grade acupuncture gold microneedles as a spatial-barcoding array patch with high-performance DNA recognition interfaces. These interfaces first utilize nanoparticles surface deposition to increase the density of DNA probes. A freeze-thawing method is further developed to regulate interfaced DNA probes with laterally uniform distribution and vertically upright conformation, mainly by loose stretching and local concentration of these DNA probes. It increased mRNA capture with 250% and speeded up reaction (from 60 min to 15 min), compared to that without freeze-thawing surface engineering. Using this method, we explored in vivo spatiotemporal transcriptome dynamics during skin wound healing of individual mice. We found four distinct expression changes of transcripts, including continuous upregulation, continuous downregulation, initial upregulation then downregulation, and initial downregulation then upregulation. Our aivST method advances the field from sparse tissue section sampling toward in vivo measurements of biological stereotypy and variability in individual living animals.

## Introduction

Spatial omics enables the revelation of intercellular communication mechanisms at the molecular interaction level. In particular, skin wound healing represents a highly coordinated and dynamic biological process^1,2^, including three distinct yet overlapping phases, inflammatory response, cellular proliferation, and tissue remodeling^3^. The precise regulation of this process relies on spatiotemporally ordered interactions among multiple cell types and cascade activation of molecular signaling. The timely initiation, efficient execution, and orderly transition of each phase are critical to wound repair^4^. Each phase involves specifically orchestrated events such as cell recruitment, differentiation, and extracellular matrix reorganization. It is well known that the spatial distribution of diverse cell types is a fundamental determinant of tissue function during skin wound repair^5,6^. Therefore, tracking spatiotemporal distribution of the transcriptomic throughout all stages is crucial for a comprehensive understanding skin wound repair mechanism.

Several approaches have been developed to address the classical limitation of resolving cellular heterogeneity while preserving spatial information, allowing researchers to visualize the spatial distribution of specific genes on tissue slices with high throughput^7^. However, current methods still rely on the analysis of tissue slices rather than living animals, not longitudinal *in vivo* monitoring^8-13^, leading to two major drawbacks. First, the sectioning process compromises cellular integrity and results in partial loss of RNA content. Second, this approach can only capture a static snapshot of biological activity at a single time point within the same sample, failing to reflect transcriptomic dynamic changes within same living animals. What’s more, the capture efficiency is constrained by the low efficiency of the surface reaction. To address these limitations, microneedles (MN), as a novel minimally invasive tool, offers an approach for repeated or continuous sampling, enabling precise collection of biomolecules *in vivo*^14,15^. Therefore, achieving *in vivo* spatial transcriptomic analysis remains unresolved challenge.

To overcome these challenges, we developed acupuncture-extracted in vivo spatiotemporal transcriptomics (aivST), a method engineers clinical-grade acupuncture gold microneedles (MNs) as a spatial-barcoding array patch with high interface capture ability. It uses both nanoparticles surface deposition and DNA freeze-thawing surface engineering to construct a high-performance molecule recognition interface. The well-known surface deposition increases the modified areas and numbers of DNA capture probes^16^. Our previous freeze-thawing DNA assembly on gold surfaces^17^ is extended to gold MNs. This process mainly drives loose stretching and local concentration of these DNA probes, thus regulating them with laterally uniform distribution and vertically upright conformation. The finite element analysis is also performed to prove that freezing-driven confinement transforms DNA transfer from a canonical diffusion-limited process into a surface-confined manner. Using total RNA as inputs, this freeze-thawing surface engineering achieved about 250% increase of mRNA capture and speeded up the reaction from 60 min to 15 min. Then different DNA barcode-functionalized MNs were used to construct the spatial-barcoding patch based on a customized 3D-printed microwell array.

As a proof of concept, we fabricated a 3×3 MN patch to monitor in vivo spatiotemporal transcriptome dynamics during skin wound healing of individual mice. The *in vivo* extraction of mRNA molecules were collected and transformed as cDNA by reverse transcription, followed by library construction, PCR amplification and sequencing analysis. We found four different expression changes of transcripts during the three distinct phases of wound healing (inflammation, proliferation, and tissue remodeling). They included continuous upregulation, continuous downregulation, initial upregulation followed by a downregulation, and initial downregulation followed by an upregulation. We also mapped the distinct transcriptome landscapes at multiple spatial locations proximal to the wound depending on different distances. Using aiVST, we directly explored spatiotemporal transcriptomics in living animals, which were impossible by formerly existing methods.

## Results

### Design of aivST-seq

As shown in Scheme 1a, current spatial transcriptome sequencing strategies are limited to tissue sections, failing to directly detect individual living tissues and living animals. Our aivST method addresses this limitation by engineering clinical-grade acupuncture gold microneedles (MNs) with a set of surface designs (Scheme 1b). The first one is engineering gold nanoparticles (AuNPs) coating via a single deposition step, forming a high-quality surface with nanostructured morphology. This AuNP-coated microneedle (AuNP-MN) both maintains the high sharpness for tissue penetration and increases the surface area for high-density immobilization of DNA capture probes by the robust Au-S chemistry. Thus, it can capture more target molecules (e.g., mRNA molecules) compared to the bare microneedle (bare-MN). Additionally, the electrodeposition results in a strong adhesion to the microneedle substrate due to the local galvanic reduction of gold ions^18^, which protects the AuNP-MN against disruptive mechanical forces such as tissue insertion.

**Scheme 1.**
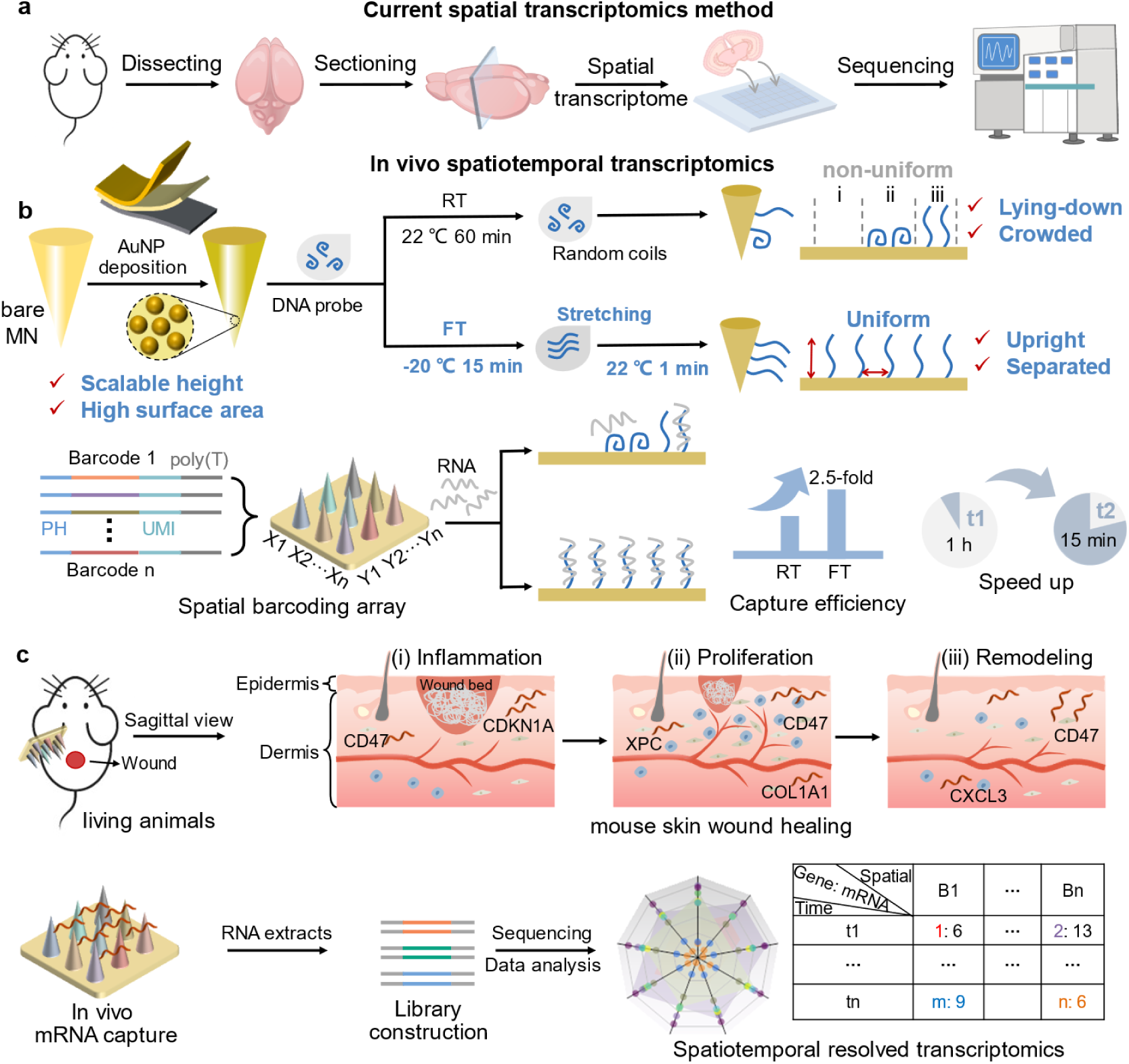
| Spatiotemporal transcriptomic profiling of skin wound healing using MN patch. (a) Current spatial transcriptome method. (b) *In vivo* spatiotemporal transcriptomics analysis, including the MN array fabrication, and state diagrams of DNA immobilization after adding salt at room temperature (RT) and through freeze-thawing (FT). (c) Schematic illustration showing work flow of AuNP-MN patch biodetection involving *in vivo* sampling and transcriptome analysis in skin wound healing.

The second one is engineering high-performance DNA recognition solid-liquid interface by a freeze-thawing process^17^. In the traditional (room temperature, RT) DNA immobilization on gold surfaces, three scenarios are presented (Scheme 1b, RT): i) bare surface without DNA probe, ii) DNA probe with a lying-down conformation, and iii) a close-packed DNA probe layer with an upright conformation. These three non-uniform probe settings all limit capture efficiency of mRNA molecules. In contrast, our developed freeze-thawing strategy can loosely align DNA with a more extended conformation rather than random coil conformation (Scheme 1b, FT), and concentrate these oligos within an interfacial liquid layer for a surface-confined reaction instead of a diffusion-mediated reaction. This ice-confinement treatment results in a high-uniform DNA recognition interface with vertically upright and laterally separated conformations. Thus, this DNA recognition interface can significantly speed up reaction and enhance the capture efficiency of mRNA molecules *in vivo*. For DNA probe design, partially complementary MN-surface probe and MN-capture probe, are used to capture mRNA molecules. The MN-surface probe contains a thiol group at 3’ termini for covalent immobilization on the gold surface (Supplementary Fig. 1a). The MN-capture probe consists of five functional regions, including a PCR handle (hybridized with MN-surface probe), an anchor region, a spatial barcode region, a unique molecular identifier (UMI) for digital counting, and a poly(T) region for binding to the 3’ poly(A) of mRNA. A strand displacement primer (c-anchor) is designed to bind to this anchor region. It can initiate DNA polymerase-mediated displacement reaction to release the complex of MN-capture probe and mRNA from the MN surface. The collected mRNA are then transformed as cDNA by reverse transcription, followed by library construction, PCR amplification and sequencing analysis.

The third engineering is constructing the DNA spatial barcode-functionalized MN array patch. As a proof of concept, we design a 3×3 MN patch based on a customized 3D-printed microwell array (Scheme 1b and Supplementary Fig. 1b, c). Different DNA barcode-functionalized MNs are trimmed with uniform height and then immobilized in the microwell array with given spatial locations. Notably, the size and length of the MN determines the spatial resolution and tissue depth for *in vivo* omics profiling, respectively. The MN patch with a center-to-center distance is 2 mm, the tip length about 1 mm, base diameter approximately 150 μm. To demonstrate the performance of our method, the mice with skin wound healing are detected. It is well known that skin wound healing is a highly coordinated biological process with three distinct phases, inflammation, proliferation, and tissue remodeling (Scheme 1c). The alterations in biomolecules such as nucleic acids in the dermis are associated with the development of numerous skin disorders. We try to explore the dynamic spatiotemporal transcriptome of skin wound healing *in vivo* by measuring spatial-barcoding (B1 to Bn) mRNA extracts at different healing time, including days 0 (Normal), 1 (Wd1), 7 (Wd7), and 14 (Wd14).

### AuNPs-functionalized MN with high binding properties

As shown in Fig. 1a. To improve the DNA binding ability, we engineered a AuNP-MN according to precious method^19^. In brief, the bare-MN was applied to form AuNP layer by electrodeposition step, which results in a strong adhesion to the bare-MN substrate due to the local galvanic reduction of Au^3+^ ions. The AuNP-MN was subjected to surface chemistry analysis compared to bare-MN, which originally had an Au-plated layer. Analysis by scanning electron microscopy (SEM), the successful deposition of AuNPs onto the bare-MN was clearly observed, with an average particle size of approximately 168 nm. In contrast, the bare-MN exhibited a relatively smooth surface (Fig. 1b and Supplementary Fig. 2a).

**Figure. 1.**
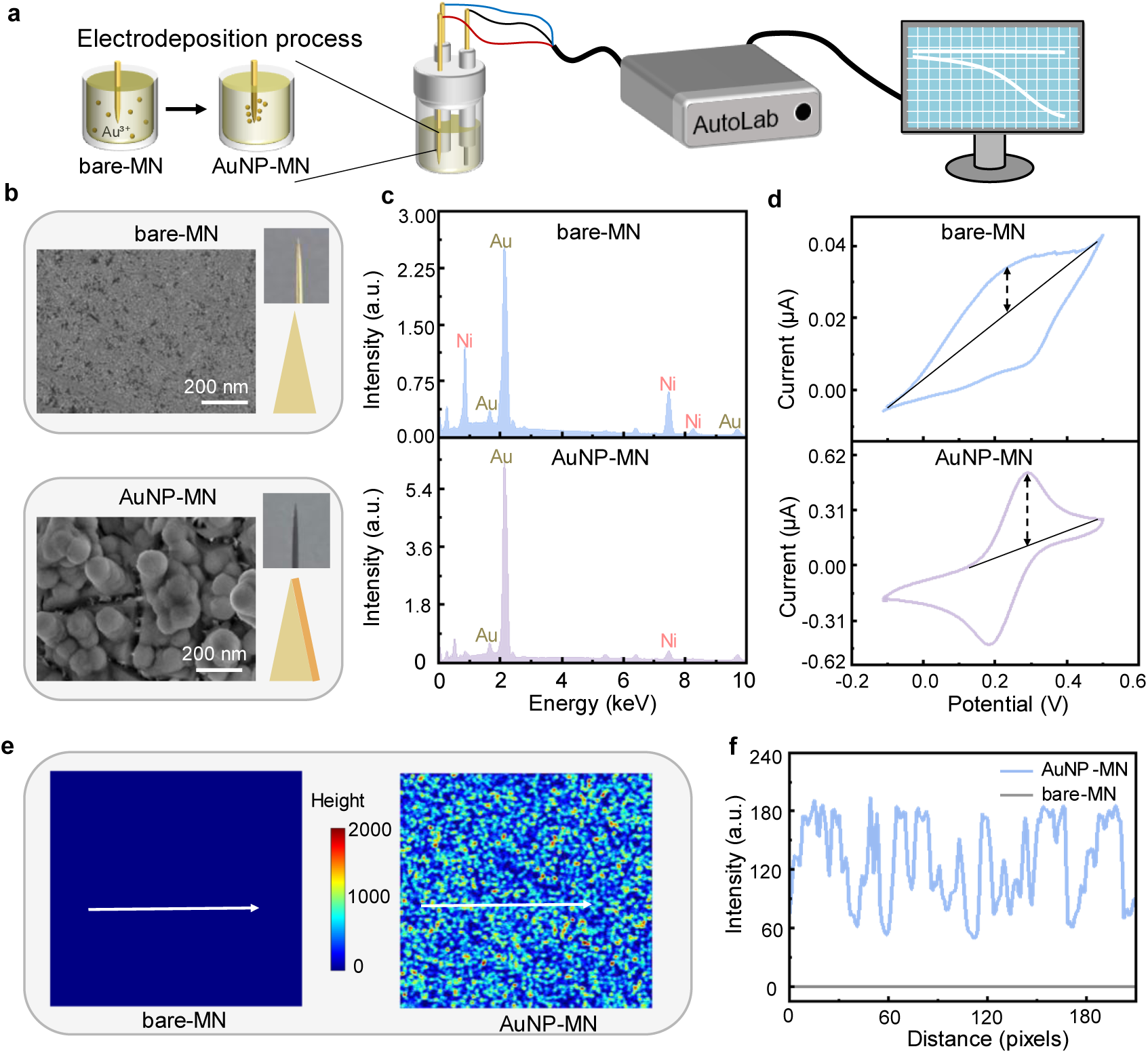
| Characterization of DNA binding ability on MN. (a) Schematic illustration of electrodeposition process. (b) The SEM image of bare-MN and AuNP-MN. Scale bar: 200 nm. (c) The EDS analysis of bare-MN and AuNP-MN. (d) Before and after AuNPs deposition, cyclic voltammetry (CV) curve of MN. (e) MATLAB simulates the surface roughness of bare-MN and AuNP-MN. Blue and red represent different height (nm). (f) The roughness curve of bare-MN and AuNP-MN.

Energy-dispersive spectroscopy (EDS) shown a significant presence of nickel (Ni) presence was observed on the bare-MN. However, the AuNPs layer effectively minimized Ni-related peaks (Fig. 1c and Supplementary Fig. 2b). The mass percentage of Ni decreased clearly from 44.27% to 12.51%, and the atomic percentage of Ni decreased from 72.27% to 32.41%. However, gold mass percentage increased markedly from 55.73% to 87.49%. Eliminating the Ni exposure is crucial, because Ni and its surface oxide layer can interfere with DNA immobilization, which is necessary for molecule detection. To confirm the successful AuNPs modification, the [Fe (CN)6]^3−^/^4−^ as redox mediators, the cyclic voltammetry (CV) shown a strong redox peak on the AuNP-MN, whereas the response on the bare-MN was weaker and difficult to distinguish from the background (Fig. 1d and Supplementary Fig. 2c). What’s more, MATLAB simulates the surface roughness of AuNP-MN and bare-MN, and it can be clearly observed that AuNP-MN has more reaction sites (Fig. 1e, f). These results indicate that AuNPs provide more active sites for DNA immobilization.

### Freeze-thawing process contributes high local concentration

To control the DNA conformation, we employed a freezing process to maintain it in an upright state (Fig. 2a). The Cy5-labeled MN-surface (MN-surface-Cy5) probe was applied to confirm the successful functionalization of the MNs. Confocal fluorescence microscopy imaging revealed that freeze-thawing-treated AuNP-MN (FT-AuNP-MN) probes exhibited strong fluorescence with a uniform distribution (Fig. 2b, c). In contrast, the room temperature-treated AuNP-MN (RT-AuNP-MN) displayed weak and scattered fluorescence, and bare-MN exhibited weak fluorescence (Supplementary Fig. 2d), suggesting a more upright conformation with reduced surface-induced quenching^20^.

**Figure. 2.**
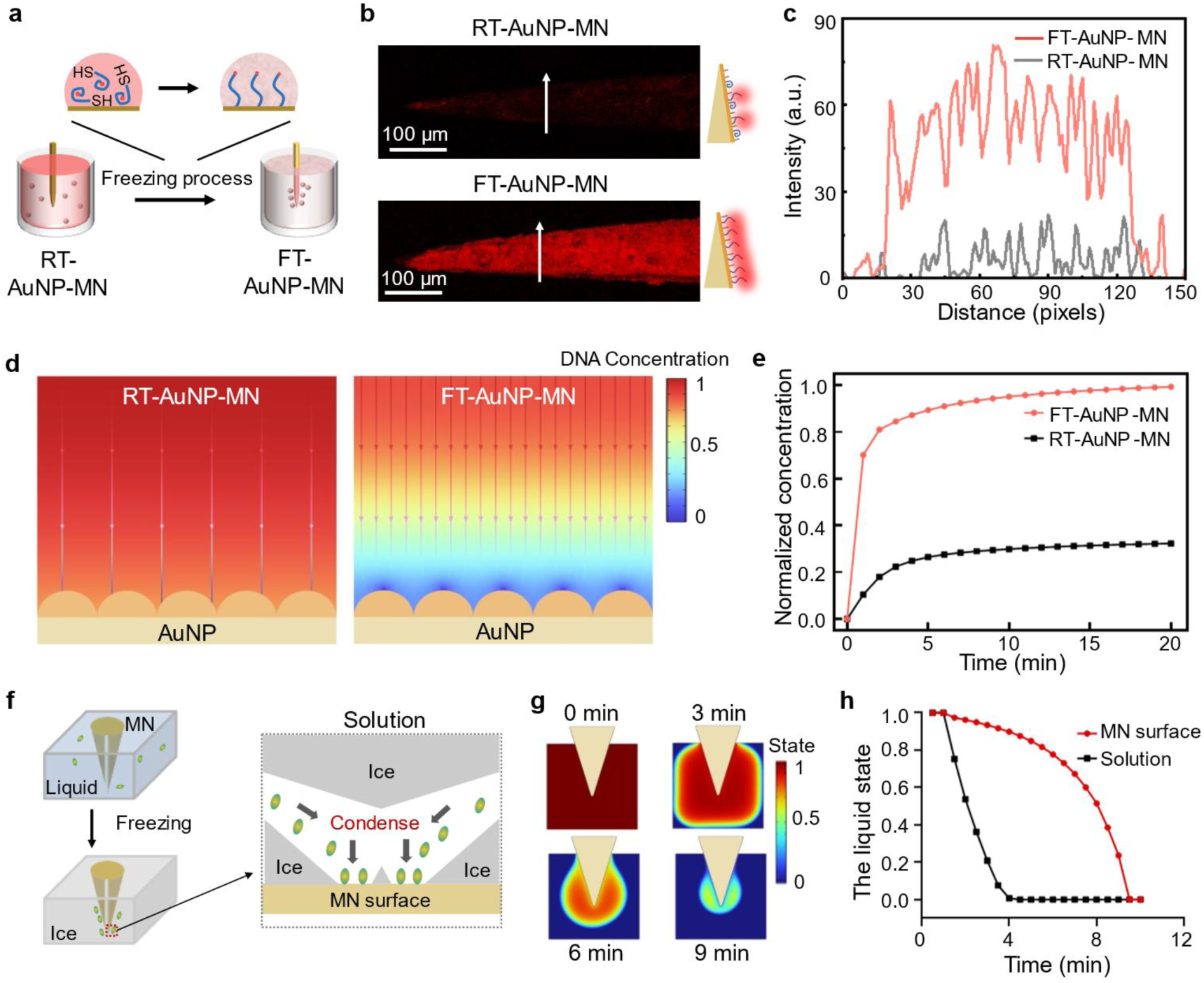
| Characterization of freeze-thawing process on MN. (a) Schematic illustration of freezing process. (b) The fluorescence image of RT-AuNP-MN and FT-AuNP-MN. Scale bar: 100 μm. (c) The intensity curves of (b). (d) COMSOL simulated DNA concentration distributions in solution and (e) normalized surface concentrations under frozen at ambient conditions. (f) Schematic illustration of directional freezing, where molecules concentrate within a thin liquid layer between ice and the MN interface. (g) COMSOL simulation of ice formation and (h) normalized temperature under frozen condition. 0 and 1 regions denote ice and liquid state, respectively.

Additionally, we employed COMSOL Multiphysics simulations confirmed these observations (Figure 2d, e), The RT-AuNP-MN showed an increase in normalized DNA concentration from 0 to 0.32, whereas the FT-AuNP-MN exhibited an increase from 0 to 0.99. This 3.1-fold enhancement could be due to the higher binding affinity of FT-AuNP-MN for cDNA. These results indicate that the ice front robustly drives DNA to the interface, generating local concentrations far exceeding those seen in bulk solution. Therefore, the combination of the freeze-thawing process with AuNPs deposition was found to contribute to an improvement in DNA binding ability. To explain freeze-thawing phenomenon (Fig. 2f, g), the COMSOL simulations were used to analyze the freezing process (Fig. 2h), revealing that freeze-concentrated solutions were driven into close and sustained contact with the MN surface.

### FT-AuNP-MN contributes high capture efficiency in vitro

To further compare the differences in RNA capture performance among the freeze-thawing assembled AuNP-MN (FT-AuNP-MN), room temperature-treated AuNP-MN (RT-AuNP-MN), and the bare-MN, we employed the TRIzol method to extract total RNA from MCF-7 cells. (Fig. 3a). The correlation analysis indicated the gene and UMI distributions between the two replicates were highly consistent in FT-AuNP-MN (*R* = 0.999), and bare-MN and RT-AuNP-MN exhibited higher correlation with each other (Fig. 3b, Supplementary Fig. 3a), this correlation further confirms the superior RNA capture ability of FT-AuNP-MN. Interestingly, we observed different numbers in three groups. The bare-MN with 368,604.67 UMIs ± 75,560.33 and RT-AuNP-MN with 424,316.67 UMIs ± 105,417.46 on the MN. FT-AuNP-MN exhibited stronger capture capability than the other two groups, with 975,680.33 UMIs ± 121,288.16 (Fig. 3c). The 2.5-fold enhancement in capture efficiency is attributed to the high density and upright conformation of the FT-AuNP-MN. We further analyzed the proportions of protein-coding and non-coding RNAs, revealing that FT-AuNP-MN captured a substantial fraction of both types compared to the other two methods (Fig. 3d). The *ACTB, CCND1*, and *ESR1* gene, as the marker gene of MCF-7 cell. Raw reads count and UMI number were detected in MCF-7 cell (Fig. 3e and Supplementary Fig. 3b, c), these results indicated that UMI counting plays a crucial role in eliminating PCR bias^21,22^. We compared the capture efficiency of the three methods for highly expressed genes in MCF-7 cells, demonstrated superior capability in detecting *ACTB*, *CCND1* and *ESR1* compared to the other methods. To further compare these methods, we used Venn diagram to assess the specificity of each approach. As shown in Fig. 3f, FT-AuNP-MN detected 23.7% more genes than conventional methods, with 4,177 additional genes compared to the bare-MN and 3,847 genes more than RT-AuNP-MN. Moreover, FT-AuNP-MN exhibited certain reproducibility in genes detection across three replicates (Fig. 3g and Supplementary Fig. 3d, e). Overall, FT-AuNP-MN provided high mRNA capture efficiency than RT-AuNP-MN and bare-MN.

**Figure. 3.**
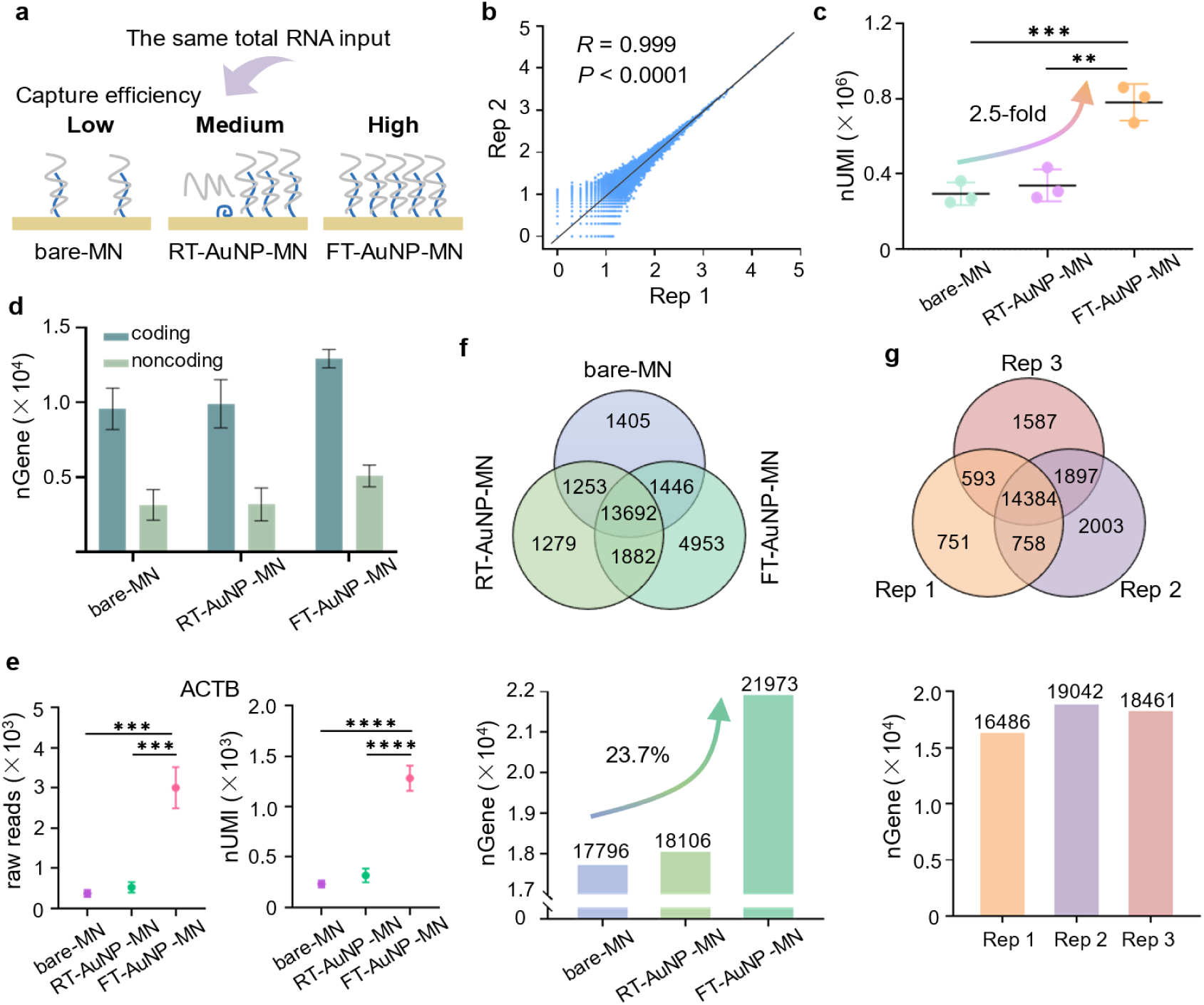
| Comparative analysis of FT-AuNP-MN, RT-AuNP-MN and bare-MN. (a) Schematic diagram of the capture efficiency for FT-AuNP-MN, RT-AuNP-MN and bare-MN. (b) Correlation of RNA level quantified by UMI numbers between two replicates (Rep) of FT-AuNP-MN. (c) Comparison of nUMI with FT-AuNP-MN, RT-AuNP-MN and bare-MN (*n* = 3 in each group), respectively; nUMI, number of UMIs. Data are presented as mean ± SD. (d) Comparison of nGene with FT-AuNP-MN, RT-AuNP-MN and bare-MN in coding and non-coding genes (*n* = 3 in each group), respectively; nGene, number of genes. Data are presented as mean ± SD. (e) The raw reads count and nUMI of *ACTB* gene detected in MCF-7 cell (*n* = 3 in each group). (f) Genes detected by FT-AuNP-MN, RT-AuNP-MN and bare-MN are plotted as Venn diagram and Bar plot, including those common to three replicates, two replicates, and those detected by only one replicate. Numbers of detected genes by 3 methods are listed. The bottom shows the number of genes detected across different of methods. (g) Genes detected in three replicates of FT-AuNP-MN. Numbers of detected genes are listed. The diagram at the bottom is the number of genes detected in three replicates.

### Fabrication and characterization of the MN patch

To validate the spatial resolution of the MN patch, as shown in Fig. 4a, we fabricated a 3×3 MN patch, the patch with a center-to-center distance is 2 mm, the tip length about 1 mm, base diameter approximately 150 μm (Fig. 4b, c). Micro-compression tests were performed to determine whether the patch exhibited sufficient strength to penetrate the epidermis and reach the dermis^23,24^. The average mechanical strength was 8.396 ± 0.871 N/MN (Fig. 4d, top), which is sufficiently strong to puncture the skin without causing the MN patch to mechanically yield. To further verify the strain of the single MN (Fig. 4d, below), we performed a finite element analysis (FEA) using COMSOL Multiphysics^25^. The simulation results illustrated stress distribution as the MN was inserted into the skin. The Equation as follow:

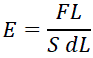

provides a theoretical proof for the MN’s penetration capability, where *E*, *F*, *S* and *L* represent the Young’s modulus, actual strain, tip area of the MN, and MN displacement, respectively. In our study, the experiment was conducted under conditions of *F* = 8 N, *L* = 2×10^-6^ m, and *S* = 7.8×10^-9^ m. The theoretical analysis indicated that *dL* approaches 0 and *E* approaches infinity within the time frame required for the MN to pierce the skin, corroborating its penetration ability.

**Figure. 4.**
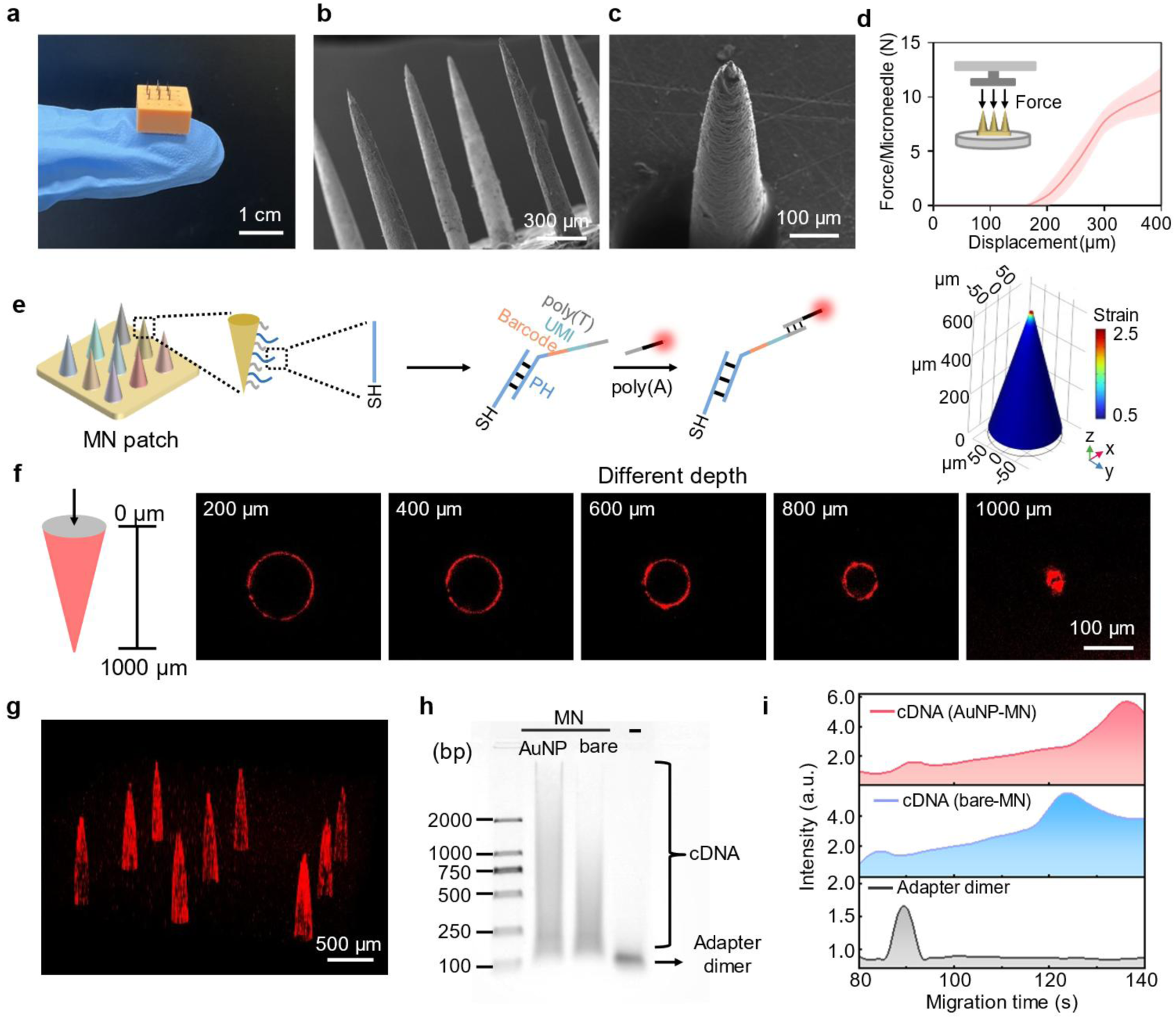
| Verification of the capturing ability of the MN patch. (a) Optical Images of the MN. Scale bar: 1 cm (b) SEM image of fabricated MN patch. Scale bar: 300 μm. (c) SEM image of singe MN. Scale bar: 100 μm. (d) Top: Mechanical behavior for the MN patch under normal compressive load and schematic illustration of experimental setup. Data are presented as mean ± SD. Below: FEA of the structure strain of the MN. (e) Schematic illustration depicting the capture efficiency. (f) The capture capabilities of different layers on the Z-axis. Scale bar: 100 μm. (g) Fluorescence imaging of MN patch. Scale bar: 500 μm. (h) Gel electrophoresis demonstrating the capture ability. (i) The length distribution of cDNA and adapter dimer.

Furthermore, to evaluate the capture efficiency, as illustrated in Fig. 4e, we employed Cy5-labeled oligo d(A) probes for verification. The SH-DNA was immobilized on the MN patch, and poly(T) probes with capture functionality hybridized with it. After a washing step, the Cy5-labeled poly(A) probes were captured, and the capture efficiency was assessed using a confocal fluorescence microscope. These results, as shown in Fig. 4f and Supplementary Fig. 4a, demonstrated that MN patch exhibit consistent capture efficiency at different depths. indicating that the MN height does not affect their capture efficiency. Furthermore, the confocal fluorescence microscope imaging revealed that MN patch exhibit consistent capture efficiency (Fig. 4g), ensuring reproducibility and reliability.

Finally, we validated the capture efficiency of the AuNP-MN patch in vitro. The total RNA extracted using the TRIzol method was characterized, after reverse transcription and template switching to synthesize cDNA, then collect cDNA via strand displacement polymerization, followed by PCR amplification. As shown in Fig. 4h and Supplementary Fig. 4b, distinct cDNA bands were observed in the RNA-containing group, whereas the bare-MN patch only exhibited adapter dimers without cDNA products. The Q-sep analysis further confirmed the AuNP-MN patch shown a prominent cDNA peak near 1,000 bp (Fig. 4i), indicating successful amplification of captured mRNAs. In contrast, the bare-MN patch showed only a peak at approximately 120 bp, corresponding to adapter dimers. These results demonstrate that the AuNP-MN patch enables effectively capture transcriptome in vitro.

### Dissecting differences of mouse skin wound healing using MN patch in vivo

Skin wound regeneration process is a regulated and complex process involving initiation, progression, and termination. To establish a mouse skin injury model, we created wounds on the dorsal skin and monitored the healing process by photography (Supplementary Fig. 5a, b), which clearly showed a gradual reduction in wound size. Before applying the MN patch for *in vivo* monitoring, the biophysical properties were systematically evaluated, including (i) the mechanical strength required for dermal penetration and (ii) its biocompatibility and safety.

To further validate penetration, Hematoxylin and Eosin (H&E) staining was conducted, confirming that the MN patch penetrated the stratum corneum and perforated into the dermal layer, with a penetration depth of approximately 180-200 μm (Supplementary Fig. 5c). Compared with conventional blood collection methods, the MN patch approach demonstrated a minimally invasive nature. Moreover, as shown in Supplementary Fig. 5d, H&E staining was also employed to assess systemic toxicity in mice, and no adverse effects were observed, indicating a favorable safety profile of the MN patch. These results demonstrate that the MN patch material had no notable adverse effects.

Furthermore, we performed a detailed analysis of transcriptional dynamic changes during mouse skin wound healing. *In vivo* sampling was conducted at days 0, 1, 7, and 14 after skin injury. (*n* = 3 in each group), followed by library construction and sequencing (Fig. 5a). Inspired by the temporal dynamics of gene expression during skin regeneration^26^, we categorized the process into four distinct modules, Module 1 (M1), Module 2 (M2), Module 3 (M3), and Module 4 (M4). Stage-specific sampling conducted throughout mouse skin repair enabled the visualization of corresponding gene expression patterns, as shown in the Fig. 5b. The *MEST*, *CXCL12* and *COL1A1* as key genes in M1, displayed a sustained upregulation throughout the repair process until complete wound closure. In contrast, M2-associated genes including *CDKN1A*, *EIF4A1* and *TRA2B* exhibited a continuous downregulation until healing was completed. Unlike M1 and M2, the expression patterns of M3 and M4 were non-monotonic. M3 signature genes (*CXCL1, CXCL2* and *HSPB1*) exhibited a characteristic pattern of initial upregulation followed by a subsequent decline, whereas M4 genes showed the opposite trend, with an earlier decrease followed by an increase. Representative genes of M4, such as *ELOVL6*, *SMAD1* and *LOR*, consistently followed this expression pattern. These results indicate that aivST-seq is capable of tracking dynamic transcriptional alterations during murine skin injury.

**Figure. 5.**
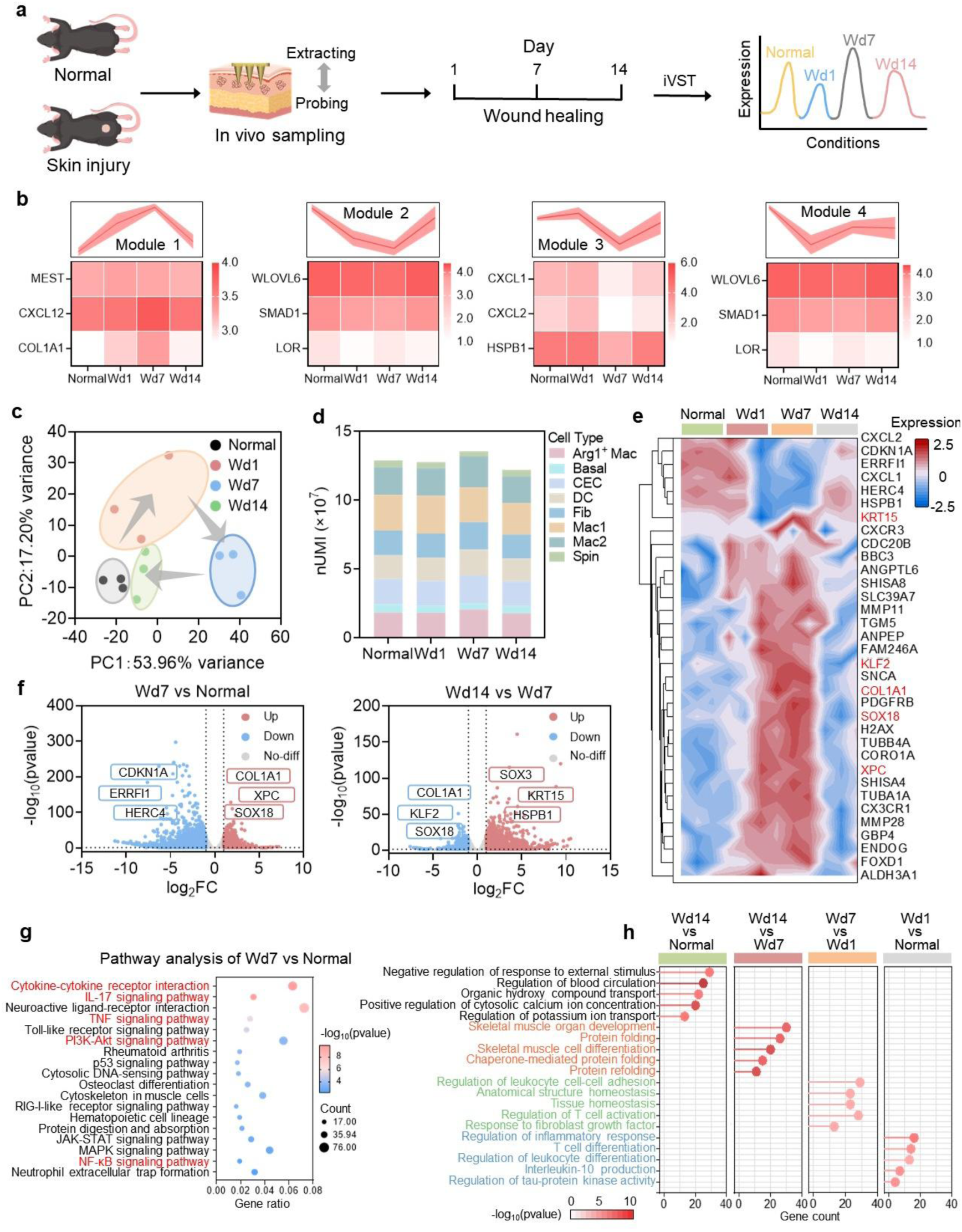
| The difference dissection of mouse skin wound healing. (a) Schematic representation of the study design employed for sampling in the skin wound healing. (b) The heatmap displays the normalized expression levels of various genes in different states (M1 to M4) and at different time points (Normal to Wd14), *n* = 3 in each group. Data are presented as mean ± SD. (c) Principal Component Analysis (PCA) analysis of samples from Normal, Wd1, Wd7 and Wd14. PC1 and PC2 represent the first and second principal components, respectively. Each point represents a sample, with colors indicating different conditions. (d) Bar plot showing nUMI for different cell types across conditions. (e) Heatmap showing clustering of normalized gene expression analysis across different conditions (n = 3 in each group). The horizontal axis represents different conditions, and the vertical axis represents gene clusters. Genes within the same branch exhibit greater similarity. Different colors represent relative gene expression levels, with red indicating high expression and blue indicating low expression. Data are presented as mean ± SD. (f) Volcano plot showing differentially expressed genes between Wd7 vs Normal and Wd14 vs Wd7. Red and blue dots represent significantly upregulated and downregulated genes, gray dots represent genes with no significant difference. Genes with a false discovery rate-corrected *P* < 0.05 and relevant genes are highlighted. (g) Kyoto Encyclopedia of Genes and Genomes (KEGG) pathway enrichment analysis of Wd7 vs Wd1, with selection criteria of |log2 FC| ≥ 1.00 and P ≤ 0.05 (*n* = 3 in each group). The vertical axis represents an enriched pathway, and the horizontal axis represents gene ratio, indicates the proportion of differentially expressed genes that are enriched in a given pathway relative to the total number of genes within that pathway. Larger value indicates a higher enrichment level of differentially expressed genes in the pathway. (h) Top 5 Gene Ontology (GO) analysis in four conditions. The horizontal axis represents gene count, and the vertical axis represents enriched biological process. Gene with selection criteria of |log2 FC| ≥ 1.00 and *P* ≤ 0.01 (*n* = 3 in each group) were used for GO analysis.

In addition, we further dissected the dynamics process of wound healing. PCA was then applied to assess similarities and differences among samples from different conditions (Fig. 5c). Samples from the same time point clustered tightly, reflecting shared transcriptional features within each condition. However, Wd7 samples showed clearly separation from the other groups, indicating a key transition point of wound healing at Wd7, characterized by peak repair activity with extracellular matrix (ECM) deposition and cell proliferation, along with pronounced transcriptomic dynamic changes. Strikingly, the samples projected onto the PCA plot formed a clear, continuous looped trajectory over time. This closed trajectory originated from the uninjured state (Normal), progressed through distinct intermediate stages, and ultimately returned to a state that closely clustered with the starting point by the end of the time course. Therefore, the PCA loop serves as a powerful visual representation of the completeness and dynamic regulation of the skin wound healing process.

Additionally, as shown in Fig. 5d. We classified cells into eight major types based on gene expression characteristics^26^, including arginase 1 positive macrophage cell (Arg1⁺ Mac), basal cell (Basal), capillary endothelial cell (CEC), dendritic cell (DC), fibroblast cell (Fib), type 1 macrophage cell (Mac1), type 2 macrophage cell (Mac2), and spinose cell (Spin). Arg1⁺ Mac is Mac expressing Arg, typically associated with anti-inflammatory responses, tissue repair, and immune regulation, which play a key role in wound healing, tumor microenvironment modulation, and certain autoimmune diseases. The normalized expression levels of Arg1⁺ Mac cells were 1.83 in the Normal group and increased slightly to 2.05 in the Wd7 group, but returned to the baseline level of 1.79 in the Wd14 group. These results suggest that the wound had largely healed at Wd14, which consistent with this observation, other cell types showed similar expression changes, which demonstrating that Wd7 as the peak phase of wound repair activity. We performed clustering analysis of key genes across four conditions, clearly illustrating stage-specific expression changes (Fig. 5e). Genes associated with the M3 module exhibited a continuous upregulation followed by a decline as wound repair progressed. In contrast, those corresponding to the M1 module, showed a sustained upregulation. The clustering further revealed that M3 module genes grouped within the same branch, indicating higher similarity among them, and similar patterns were observed for the other modules. As the peak phase of skin wound repair, the Wd7 condition exhibited gene expression levels that were markedly distinct from those at other conditions. As shown in Fig. 5f and Supplementary Fig. 6a differential gene expression analysis (|log2 FC | ≥ 1.00; *P* ≤ 0.05) were compared between Wd7 versus Normal group (Wd7 vs Normal) and Wd14 vs Wd7 group. We identified 978 upregulated and 2,191 downregulated genes in the Wd7 vs Normal group, while there were 1,370 upregulated and 713 downregulated genes in the Wd14 vs Wd7 group. and 57 upregulated and 622 downregulated genes in the Wd14 vs Normal group. Additionally, there are 176 upregulated and 260 downregulated genes in the Wd1 vs Normal group, and 141 upregulated and 967 downregulated genes in the Wd7 vs Wd1 group. These results demonstrate that different gene expression across different conditions. Furthermore. We observed that several key genes, including *COL1A1*, *XPC* and *SOX18*, were upregulated in the Wd7 vs Normal group, whereas they exhibited the opposite trend in the Wd14 vs Wd7 group, which the wound had largely healed at Wd14. Similarly, the *CDKN1A* gene provides an early DNA repair window and later downregulated to release proliferation inhibition. Consistently, in the Wd7 vs Normal group, *CDKN1A* gene expression was decreased, allowing skin cells to reenter proliferation and migration program.

Furthermore, to elucidate the biological functions of differentially expressed genes (DEGs), we performed KEGG pathway enrichment analysis (Fig. 5g and Supplementary Fig. 6b). They were found to be implicated in several key signaling pathways, including TNF^27^, IL-17^28^, PI3K-AKT^29^, and NF-κB^30^ pathway, which broadly involved in various physiological and pathological processes such as inflammation, cell proliferation, differentiation, apoptosis, and immune regulation. The Neuroactive ligand-receptor interaction signaling pathway may be involved in the regulatory mechanism of wound repair, these results provide a novel view that advances molecular diagnostics for precision medicine. Additionally, as shown in Fig. 5h and Supplementary Fig. 7, the GO enrichment analysis revealing biological processes related to wound repair, including regulation of inflammatory response, tissue homeostasis and regulation of T cell activation. These results suggest that several key pathways work in coordination to achieve inflammatory skin wound repair. What’s more, these results offer insights into the molecular mechanisms underlying skin wound repair in mice.

### Deciphering mouse skin wound healing spatiotemporal heterogeneity with aivST-seq

To investigate the spatiotemporal dynamic changes of skin wound repair, we performed *in vivo* sampling using a DNA-barcoded MN patch at different stages of skin injury in the same mice (Fig. 6a). By analyzing the expression patterns of different cell types associated with specific DNA barcodes, we observed distinct spatiotemporal specificity at the wound repair stage (Fig. 6b), different cell types showed different expression pattern in different barcodes (Bn). During the Normal and Wd14 stages, different cell types exhibited almost identical expression patterns across various barcodes, indicating that the Wd14 wound had already healed. Wd7, as the peak healing phase, different cell types exhibit different expression patterns compared to other stages, especially for CEC and Mac2 (red arrow). The closer to the wound, the higher the expression level. At Wd14, it was clearly observed that the Mac2 cell type (likely referring to macrophages) distal to the wound showed low expression, indicating that the area had completely healed. These results indicate that different expression levels of each cell type in different spatial positions.

**Figure 6.**
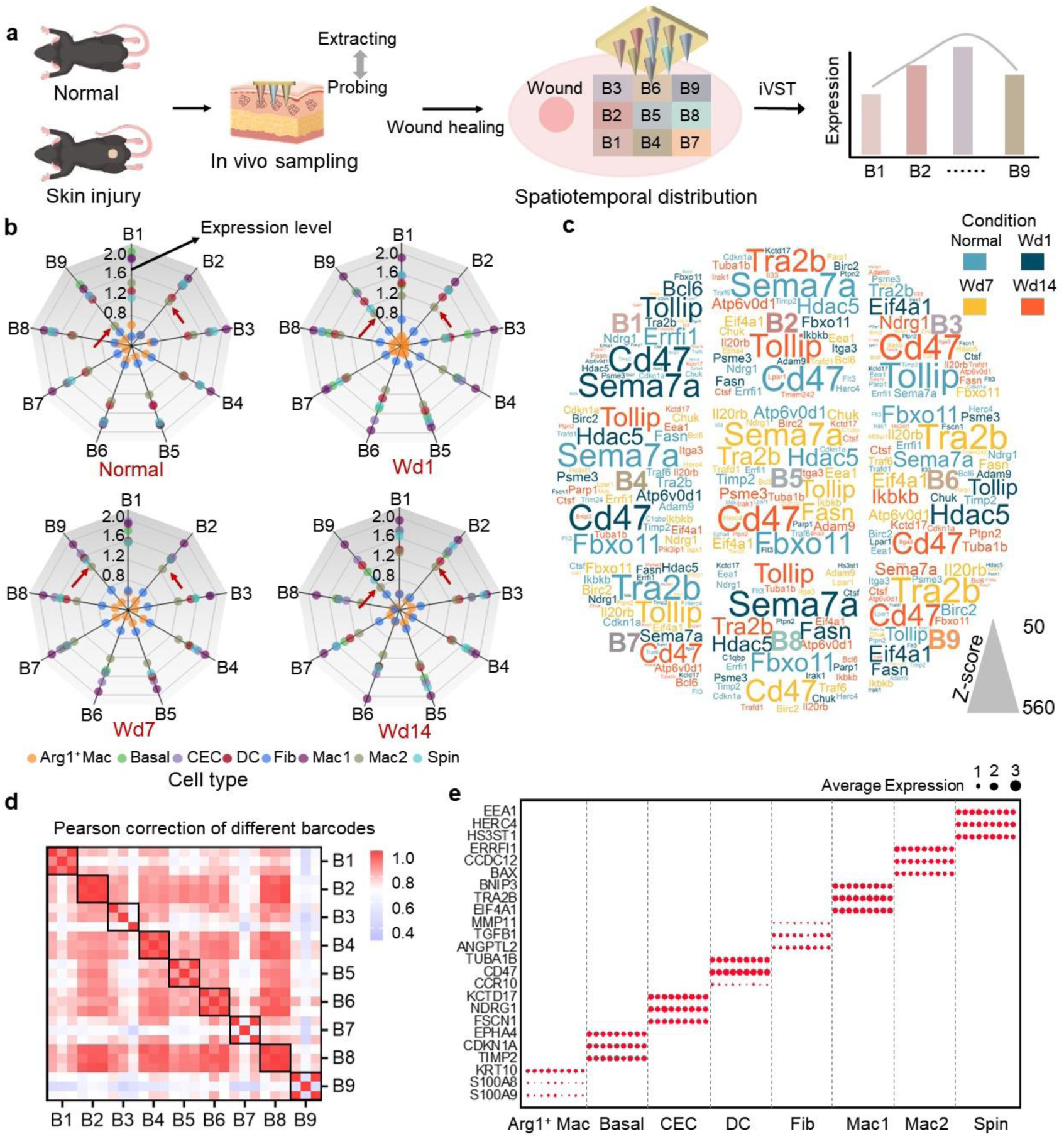
Spatiotemporal dissection of mouse skin wound healing. (a) Schematic representation of the study design employed for sampling in the skin wound healing at different spatial locations. (b) Reader chart displays the average gene expression levels at different spatial locations and at different conditions (*n* = 3 in each group). Dots represent cell type, different colors represent various cell types, the greater the distance from the center, the higher the gene expression level. B1-B3 represent proximal, B4-B6 represent middle, B7-B9 represent distal. Data are presented as mean ± SD. (c) Word cloud plots showed spatiotemporal gene expression landscape, specifically depicting the enrichment of genes with normalized expression levels (*n* = 3 in each group). The size of the letters represents the number of expression level, and the color represents the different conditions. Data are presented as mean ± SD. (d) Heatmap showing pearson correction analysis of different barcodes in Wd7. Different colors represent correction of different barcodes (blue to red, increasingly correlative). (e) Dot plot showing the average expression level of distinctive representative gene for each cell type (*n* = 3 in each group). The bubble size represents average expression. Data are presented as mean ± SD.

To describe the differentially expressed genes, we utilized word cloud plot described in detail the gene expression level across different spatial locations and at different conditions (Fig. 6c). *CD47* gene dominated in different spatial locations, and different gene exhibited distinct spatial distribution patterns. Furthermore, we examined the spatial distribution of marker genes related to wound healing (Supplementary Fig. 8a). The *XPC* gene (the recognizer for DNA damage repair), *COL1A1* gene (the key collagen for skin repair), *KLF2* gene (key transcription factor for immune regulation) and *SOX18* gene (regulates endothelial cell differentiation) were significantly upregulated at Wd7 stage, with the higher expression levels in barcodes closest to the wound, indicating that expression level is spatially correlated with proximity to the injury wound. As the peak healing phase, we further assessed the correlation of gene expression across different barcode at Wd7 (Fig. 6d), demonstrating comparable expression levels among different barcodes, particularly adjacent barcodes. Consistent correlations were also demonstrated in other stages (Supplementary Fig. 8b). These results indicate shared cellular composition or functional states in adjacent areas. Additionally, we observed enrichment of specific cell types in different barcodes, by examining the expression distribution of representative genes for different cell types at Wd7, reflecting spatially organized cellular distribution and functional specialization (Fig. 6e). Overall, aivST-seq revealed a potential spatiotemporal trajectory of skin wound healing in mice.

## Discussion

Spatial transcriptomic enables precise mapping of gene expression patterns across distinct regions within tissue slices, playing a vital role in elucidating spatial heterogeneity. However, this approach currently relies on the analysis of tissue sections, which limits its ability to capture transcriptional changes in response to temporal variations and microenvironmental dynamic changes in living animals. This constraint is particularly evident in complex and multi-stage biological processes, such as skin wound healing. During the preparation of this manuscript, a related study focusing on spatiotemporal lipidomics in live tissues was published online^31^. However, the technical approach employed in that work still exhibits certain limitations, primarily characterized by low interface capture efficiency and insufficient detection sensitivity. To address this limitation, aivST-seq technology as an innovative method for monitoring gene expression *in vivo*, it allows longitudinally transcriptomic profiling of the same living animals at multiple time points. Several strategies could have contributed to aivST-seq high capture efficiency in gene detection First, the properties of AuNPs coating have increased the transcriptome capture efficiency, forming a high-quality surface with nanostructured morphology. Second, we integrated a freeze-thawing technique to achieve oriented assembly of DNA probes. This process mainly drives loose stretching and local concentration of these DNA probes, maintaining them in an upright conformation. The combination of these two strategies had increased the identification efficiency by 250% and 23.7% more genes were detected. Additionally, we performed *in vivo* sampling of mouse skin wounds under different conditions, and identified *KLF2*, *COL1A1*, *XPC* and *SOX18* as the primary cytokines/chemokines produced by keratinocytes in acute mouse wounds, and the related gene pathways, including TNF, IL-17, PI3K-AKT, and NF-κB, were identified. These genes have been recognized as critical regulators in skin wound healing. Furthermore, we dissected the spatiotemporal differences of mouse skin wound healing, different gene exhibited distinct spatial distribution patterns. *CD47* gene dominated in different spatial locations. What’s more, a gradient of increasing expression in repair-related genes with closer proximity to the wound. In contrast to previous studies that primarily emphasized tissue sections, aivST-seq offers a comprehensive view of spatiotemporal landscapes *in vivo*, allowing us to explore the dynamic cellular processes and regulatory networks underlying the gene expression signatures.

The aivST-seq method provides a novel tool for spatiotemporal omics profiling *in vivo*. In principle, it is also appropriate for time-resolved dynamic transcription profiling *in vivo* at the single-animal level by examining new and old RNAs. In addition, we can even utilize antibodies to capture target molecules for spatiotemporal proteomics studies, and are capable of isolating organelles with spatiotemporal resolution, thereby enabling in-depth research at the organelle level. Beyond applications in skin regeneration, aivST-seq is a versatile platform that can be extended to investigate diverse biological processes such as aging, exercise, and other physiological or pathological models within the same individual. Furthermore, we plan to integrate aivST-seq technology with acupuncture therapy to systematically analyze the biological responses at acupuncture sites. Although our study focused on mammalian cells, the underlying design of aivST-seq could be adapted for use in other organisms. Given that Omics technology had been used in bacterial^32^ and plant cells^33^, we anticipate that aivST-seq may also be applicable in these systems.

Looking forward, aivST-seq still has room to further improve performance. The platform may be limited by endogenous nuclease activity, leading to nucleic acid degradation. Therefore, incorporating chemical modifications such as Locked Nucleic Acids (LNA) could enhance nuclease resistance^34^ and improve hybridization affinity for more robust *in vivo* applications. Higher capture efficiency and gene detection sensitivity could be achieved by generating more favorable RNA capture substrates, such as combining dendrimers, which have 64 active groups on the periphery (diameter of ∼4.5 nm), could provide nanosubstrates of considerable surface roughness over the planar surface^35,36^, increasing available surfaces for DNA modification. The current aivST-seq platform is limited to transcriptome-wide analysis. Future developments aim to enable multimodal profiling of the entire or targeted transcriptome, proteome^10,37^, epigenome^38^ and chromatin modifications^39,40^ in living animals, and we will allow dynamic tracking of molecular events *in vivo* through integration with RNA dynamics^41,42^ technology, as demonstrated by other spatial molecular profiling technologies. Furthermore, we anticipate that future refinements will significantly enhance its spatial resolution. This may involve the fabrication of nanoneedles^31,43^ to reduce tip-to-tip spacing, or the integration of microfluidic technologies to increase throughput while simultaneously minimizing space. Currently, our method lacks the capability to effectively resolve structural differences across distinct tissue layers. In the near term, we will further optimize Z-axis barcoding technology to enable deep-resolved spatiotemporal omics analysis *in vivo*.

## Acknowledgements

This work was supported by the National Natural Science Foundation of China (No. 22125404, 22522409 and 92068118), the Natural Science Basic Research Program of Shaanxi (2023-JC-JQ-13), the Innovation Capability Support Program of Shaanxi (2023-CX-TD-62). We thank Ying Hao (Instrument Analysis Center of Xi’an Jiaotong University) and Liying Liu (Biomedical Experimental Center of Xi’an Jiaotong University) for the assistance with confocal fluorescence imaging analysis; Zhi Geng (the National Innovation Platform for Industry-Education Integration of Energy Storge Technology of Xi’an Jiaotong University) for the assistance with scanning electron microscope analysis.

## Author Contributions

C.H.F., Y.X.Z. and F.C. directed the research. Y.B.G., F.C. and Y.L.Z. designed the experiments and wrote the manuscript. Y.B.G. and Y.L.Z. performed the experiments and data analysis. Y.B.G., X.Y.L., J.W.K. and K.C. created the schematics and data visualizations. Y.B.G., F.C. and Y.X.Z. wrote and revised the manuscript. All authors have given approval to the final version of the manuscript.

## Supporting Information

Materials and methods, figures for additional explanation and the DNA oligonucleotide sequences used in the experiment.

