## Supplementary Information for "DNA-barcoded microneedle array patch enabling the profiling of in vivo spatiotemporal transcriptomics"

### Materials and Methods

#### Mice

The study was carried out using C57BL/6J mice (6-8 weeks, Animal Feeding Center, Xi'an Jiaotong University Medical School. Shaanxi). All animal experiments were approved by the Biomedical Ethics Committee of Health Science Center of Xi'an Jiaotong University (2021-270). Animals were housed under specific-pathogen-free conditions at 24 °C and 35% humidity, with a 12 h light-dark cycle.

#### Cell lines and reagents

The MCF-7 cell line was obtained from Pricella. All cell cultures were maintained in a 37 °C incubator with 5% $CO_2$. The culture medium used in this study was high glucose Dulbecco's modified Eagle's medium (DMEM) supplemented with 10% fetal bovine serum and 1% penicillin-streptomycin solution (all from Gibco). Gold-plated acupuncture microneedles (MN) were purchased from Suzhou Acupuncture and Moxibustion Appliance Co. Ltd. (Suzhou, China). Poly (dimethylsiloxane) (PDMS) was obtained from Dow Corning (MI, Germany). DNA oligonucleotides were purchased from Sangon (Shanghai, China). The sequences of DNA oligonucleotides used in this work were listed in Supplementary Table 1. Bst DNA polymerase (Large fragment) and Q5 hotstart polymerase were purchased from New England Biolabs (Beijing, China). Ampure XP Beads and TruePrep DNA Library Prep Kit V2 for Illumina were purchased from Vazyme (Beijing, China). Maxima H Minus Reverse Transcriptase was purchased from Thermo Fisher Scientific (MA, USA). All solutions were prepared using DEPC-treated water.

#### Preparation of DNA-barcoded MN patch

The MN patch consists of a three-dimensionally (3D) printed mould (as the housing of the MN patch), MNs and PDMS microwell (as the reaction chamber). The MNs were made according to the previously reported literature, but slightly transformed here[S1]. To form the gold nanoparticle (AuNP) coating, the electrodeposition process was applied. First, MNs were washed with absolute ethanol and ultrapure water for 5 min each, and then placed in the deposition solution (1.2 mg $mL^{-1}$ chloroauric acid solution, 0.1 M potassium chloride and 1.5% (wt, %) hydrochloric acid). Then a pulse voltage of 0 and -0.4 V was applied to 240 scanning bands for forming AuNP layer. DNA probes were immobilized on the MNs using a previously reported freeze-thawing method, but slightly transformed here[S2]. To reduce the disulfide bond in thiolated DNA, 5 μL of 100 mM Tris (2-carboxyethyl) phosphine hydrochloride (TCEP) was mixed with 10 μL of 100 μM N-surface for 1 h at room temperature in the dark. To fabricated DNA-barcoded MN patch, we have two methods for assembling DNA. One is based on freeze-thawing process (FT), and the other is to keep it at room temperature all the time (RT). For FT, reduced N-surface probes were dropped into the freshly cleaned PDMS chamber at the concentration of 1 μM in 20 mM Tris-HCl (pH 7.4), then the patch was placed in a laboratory freezer (-20 °C) for 15 min. After thawing at room temperature (22 °C) for 1 min, the resulting patch was washed with 20 mM Tris-HCl (pH 7.4) three times. For RT, reduced N-surface probes were dropped into the freshly cleaned PDMS chamber, then the patch was placed at room temperature for 1 h. Then the patch was washed by the reaction buffer three times. To backfill the MN patch surface, both methods were incubated with 1 mM 6-mercapto-1-hexanol (MCH) solution for 1 h at room temperature. Then the 1 μM N-capture probes were hybridized with N-surface probes in hybridization buffer (20 mM Tris buffer, pH 7.4, and 12.5 mM $MgCl_2$,), which have different barcode in different chamber. The MNs were rinsed for future use.

**Characterization of MN patch**

The micro-topography of the MN patch was characterized by SEM (ZEISS, Sigma 360), the surface chemical compositions were characterized by EDS. Electrochemical determination was conducted with a AUTOLAB electrochemistry workstation (Metrohm, PGSTAT302N). Fluorescence image was conducted by confocal microscope (Leica TCS SP8). Images were collected utilizing a 10×objective, and subsequent image processing was performed using imageJ software. Mechanical properties of MNs were characterized by electronic universal tester (CMT6103). The mechanical properties of MN patch were measured by putting the MN patch on a stainless-steel plate surface. The moving speed was 1 mm $min^{-1}$ until the fracture force was recorded.

**Total RNA Extraction**

Total RNA was extracted using the TRIzol (Thermo Fisher) method. MCF-7 cell reached a maximum confluence of 90%, 1 mL of TRIzol was mixed with the cells for 5 min to allow complete disruption of protein and RNA cross-linking and release of RNA. Then, 0.2 mL of chloroform was added, and vigorously shaken for 15 min at 4 ℃ at 12,000 g. This resulted in three layers (top: RNA-containing aqueous phase, middle: DNA layer, bottom: organic phase). Carefully transfer the supernatant (aqueous phase) to a new tube, then add 0.5 mL of isopropanol and incubate for 10 min. Centrifuge the tube at 4 ℃, 12,000 g for 10 min. Total RNA precipitate forms a white gel-like pellet at the bottom of tube. Discard the supernatant, then add 1 mL of 75% ethanol to wash away the excess impurities, and centrifuge the tube at 4 ℃ for 5 min at 7, 500 g, repeat this step twice. Finally, dry the RNA precipitate for 5 min, dissolve it with DEPC water, and determine its concentration using NanoDrop spectrophotometer (Thermo Fisher Scientific).

**Comparative analysis of bare-MN, RT-AuNP-MN and FT-AuNP-MN in vitro**

To further compare the differences in RNA capture performance among the freeze-thawing-treated AuNP-MN (FT-AuNP-MN), room temperature-treated AuNP-MN (RT-AuNP-MN), and the bare-MN. For FT-AuNP-MN and bare-MN, MN-surface probes were dropped into the freshly cleaned PDMS chamber at the concentration of 1 μM in 20 mM Tris HCl (pH 7.4), then the patch was placed in a laboratory freezer (-20 ℃) for 15 min. After thawing at room temperature (22 ℃) for 1 min, the resulting patch was washed with 20 mM Tris-HCl (pH 7.4) three times. For RT-AuNP-MN, MN-surface probes were dropped into the freshly cleaned PDMS chamber, the patch was placed at room temperature for 1 h. To backfill the MNs surface, both methods were incubated with 1 mM MCH solution for 1 h at room temperature. After rinsing, they were applied for hybridizing MN-capture probes, the resulting patch was washed with 20 mM Tris-HCl three times. Then reverse transcription was conducted for 2 h at 42 ℃ using 20 μL reverse transcription mix containing 4 μL of Maxima 5×RT Buffer, 2 μL of 10 mM dNTPs, 1 μL of Maxima H Minus Reverse Transcriptase, 4 μL of 20% Ficoll

PM400, 0.5 μL of BSA (4 mg $mL^{-1}$), 2 μL of Actinomycin D (500 ng $μL^{-1}$), 0.5 μL of RNase inhibitor (10 U $μL^{-1}$), 0.5 μL of 100 μM template switch oligo 4.5 μL of DEPC-treated water and 1 μg total RNA. After the cDNA was synthesized, c-anchor probes were added, allowing for the subsequent strand-displacing polymerization reaction. The reaction was conducted for 2 h at 37 °C to extend anchor oligo until displacement the cDNA from MNs.

**Amplification and sequencing**

The resulting cDNA (20 μL) from each sample was amplified with 25 μL of 1×Q5 Hotstart ReadyMix and 5 μL of cDNA primers (10 μM). The following PCR program was used: 98 °C for 3 min; five cycles of 98 °C for 20 s, 63 °C for 20 s and 72 °C for 1 min; fifteen cycles of 98 °C for 20 s, 67 °C for 30 s and 72 °C for 1 min and a final incubation at 72 °C for 5 min. PCR products were purified using the VAHTS DNA Clean Beads at a 0.6×bead per sample ratio. Final concentrations were quantified with a Qubit dsDNA assay kit. A total of 50 ng of cDNA was fragmented with a TruePrep Homo-N7 DNA Library Prep kit following the manufacturer's instructions, and the resulting library was quality controlled and and sequenced on the NovaSeq Xplus platform (Illumina).

**Spatiotemporal dissection of mouse skin wound healing *in vivo***

Female C57BL/6J mice were randomly divided into four groups ($n$ = 3 in each group), the mouse was under isoflurane anesthesia (1.5 to 2%) with a circulating heating pad to maintain its body temperature at 37 °C. When making the wound, the use of sharp, thin and curved surgical scissors is recommended. All surgical tools should be properly sterilized before use. An initial cut should be made to allow the surgical scissors to penetrate under the skin (with hair removed). After wound created, a functioned MN patch was applied onto the mouse's lower back wound, and a Tegaderm film (3M, USA) was used to secure the device in place. the applicability of this tape for connecting skin and devices have already demonstrated by previous studies[S3, S4]. After 30 min of RNA capture, the MN patch was used for reverse transcription to synthesize cDNA, and a strand displacing polymerase subsequently was conducted for 2 h at 37 °C to extend anchor oligo until displacement the cDNA from MN patch. Purified libraries were stored at -20 °C for downstream sequencing.

**Data analysis of aivST-seq**

Fastq files were generated using an Illumina NovaSeq Xplus sequencer. First, quality control of the raw sequencing data was performed by filtering low-quality bases using a sliding window algorithm (window size 4 bp, quality threshold Q30). According to the structural characteristics of the library, the molecular labeling information was located and extracted from Read 1. The specific extraction strategy was as follows: identify the anchor area as the position point, and sequentially extract the cellular barcode sequence (8 bp) and UMI sequence (10 bp) at its 3' end. The extracted barcodes were corrected using a Hamming distance-based error correction algorithm. UMI de-duplication is performed using a directed acyclic graph-based algorithm. Then gene expression count data were extracted from the count files (.txt format) of each sample, which were generated by RNA-seq alignment and counting software.

Differential expression analysis is performed using the DESeq2 package[S5], which is specifically designed for statistical analysis of RNA-seq count data based on a negative binomial distribution model. Differentially expressed genes were screened using a dual criterion of p-value < 0.05 and |log2 FC| > 1.00. Gene Ontology (GO) enrichment analysis and Kyoto Encyclopedia of Genes and Genomes (KEGG) analysis was implemented by the ClusterProfiler R packages. All analyses were conducted in a Linux environment to ensure optimal utilization of computational resources and reproducibility of analyses.

The sequencing data generated in this study have been deposited at Gene Expression Omnibus (GEO) with accession number GSE309997.

**Statistics and reproducibility**

All statistical analyses were performed on GraphPad Prism 9.5 software and Origin 2019. All of the results are presented as mean ± SD. Statistical analysis was performed between two groups using an unpaired two-tailed student's t-test. For multiple groups, one-way analysis of variance (ANOVA) with Tukey's multiple comparisons test were applied. The differences between the experimental group and the control group were considered statistically significant at $P < 0.05$: *$P < 0.05$, **$P < 0.01$, ***$P < 0.001$ and ****$P < 0.0001$. All experiments were independently repeated at least three times. The numbers of experimental replicates (n) are indicated in the figure legends.

**Supplementary Table 1. Oligonucleotide sequences used in this manuscript.**

| Oligo name | Sequence (from 5’ to 3’) |
|---|---|
| MN-surface | GACTCGACTGGAGCGTCGTGTAGGGAAAGA-HS |
| c-anchor | GGAGACAAAGATCGG |
| MN-capture1 | TCTTTCCCTACACGACGCTCTTCCGATCTTTGTCTCCAAACATCGNN NNNNNNNNTTTTTTTTTTTTTTTTTTTTTTTTTT*T*T*T*T*V*N |
| MN-capture2 | TCTTTCCCTACACGACGCTCTTCCGATCTTTGTCTCCAACGTGATNN NNNNNNNNTTTTTTTTTTTTTTTTTTTTTTTTTT*T*T*T*T*V*N |
| MN-capture3 | TCTTTCCCTACACGACGCTCTTCCGATCTTTGTCTCCACAAGCTANN NNNNNNNNTTTTTTTTTTTTTTTTTTTTTTTTTT*T*T*T*T*V*N |
| MN-capture4 | TCTTTCCCTACACGACGCTCTTCCGATCTTTGTCTCCACCACTGTNN NNNNNNNNTTTTTTTTTTTTTTTTTTTTTTTTTT*T*T*T*T*V*N |
| MN-capture5 | TCTTTCCCTACACGACGCTCTTCCGATCTTTGTCTCCAGTGGTCAN NNNNNNNNNTTTTTTTTTTTTTTTTTTTTTTTTTT*T*T*T*T*V*N |
| MN-capture6 | TCTTTCCCTACACGACGCTCTTCCGATCTTTGTCTCCATGCCTAANN NNNNNNNNTTTTTTTTTTTTTTTTTTTTTTTTTT*T*T*T*T*V*N |
| MN-capture7 | TCTTTCCCTACACGACGCTCTTCCGATCTTTGTCTCCCAGATCTGNN NNNNNNNNTTTTTTTTTTTTTTTTTTTTTTTTTT*T*T*T*T*V*N |
| MN-capture8 | TCTTTCCCTACACGACGCTCTTCCGATCTTTGTCTCCCATCAAGTNN NNNNNNNNTTTTTTTTTTTTTTTTTTTTTTTTTT*T*T*T*T*V*N |
| MN-capture9 | TCTTTCCCTACACGACGCTCTTCCGATCTTTGTCTCCCGCTGATCNN NNNNNNNNTTTTTTTTTTTTTTTTTTTTTTTTTT*T*T*T*T*V*N |
| poly(A) | Cy5-AGACGTGTGCTCTTCCGATCTAAAAAAAAAAAAAAAAAAAA |
| MN-surface-Cy5 | Cy5-GACTCGACTGGAGCGTCGTGTAGGGAAAGA-HS |
| TSO | AAGCAGTGGTATCAACGCAGAGTACATrGrGrG |
| Seq-1 | ACACTCTTTCCCTACACGACGCTCT |
| Seq-2 | AAGCAGTGGTATCAACGCAGAG |
| i5 primer | AATGATACGGCGACCACCGAGATCTACAC[i5]ACACTCTTTCCCTAC ACGACGCTC |

Cy5- denotes a 5’ Cy5 modification, -HS denotes a thiol modification, the “N” and “V” represents a degenerate base. The letter “r” indicates that the nucleotide at this position is a ribonucleotide, * indicates phosphothioate modification.

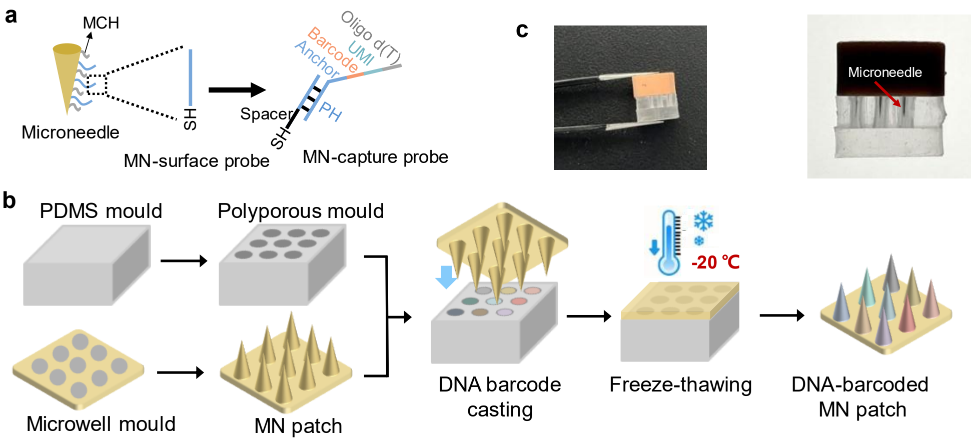

**Supplementary Fig. 1. The MN patch fabrication process.** (a) Schematic of a DNA barcode-functionalized MN, with functional domains annotated. (b) The schematic diagram of MN patch fabrication process. (c) Photograph of the MN patch at freezing stages. The red arrow indicates the MN.

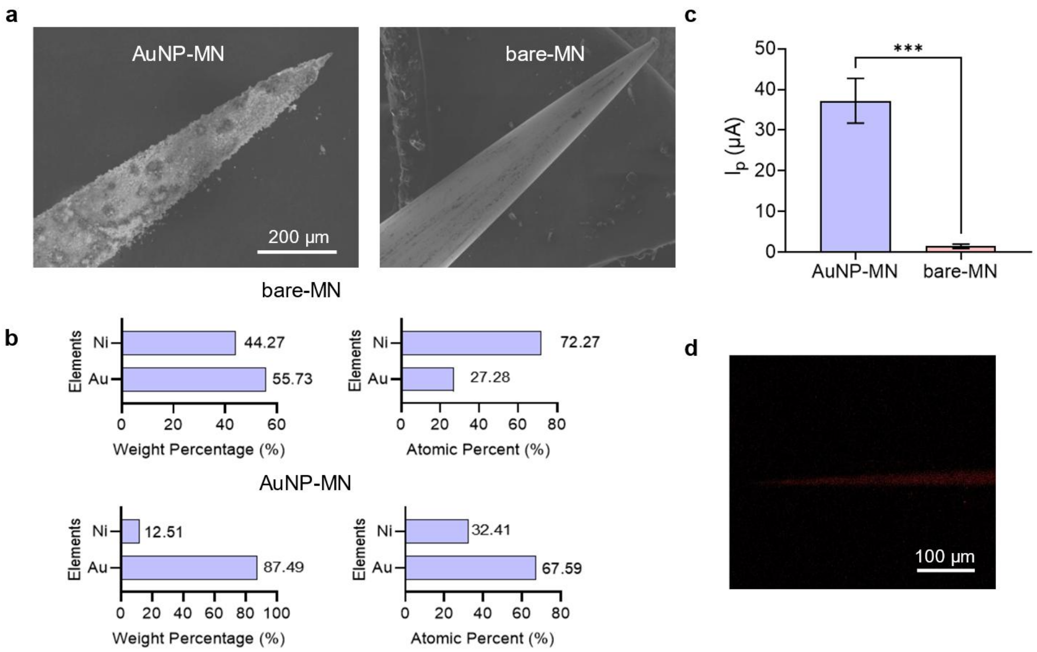

**Supplementary Fig. 2. The morphology and properties of MNs.** (a) SEM image of the bare-MN and AuNP-MN. Scale bar: 200 μm. (b) EDS mapping of (a). (c) The current vector diagram for the bare-MN and AuNP-MN ($n$ = 3 in each group). Data are presented as mean ± SD. (d) The fluorescence image of bare-MN. Scale bar: 100 μm.

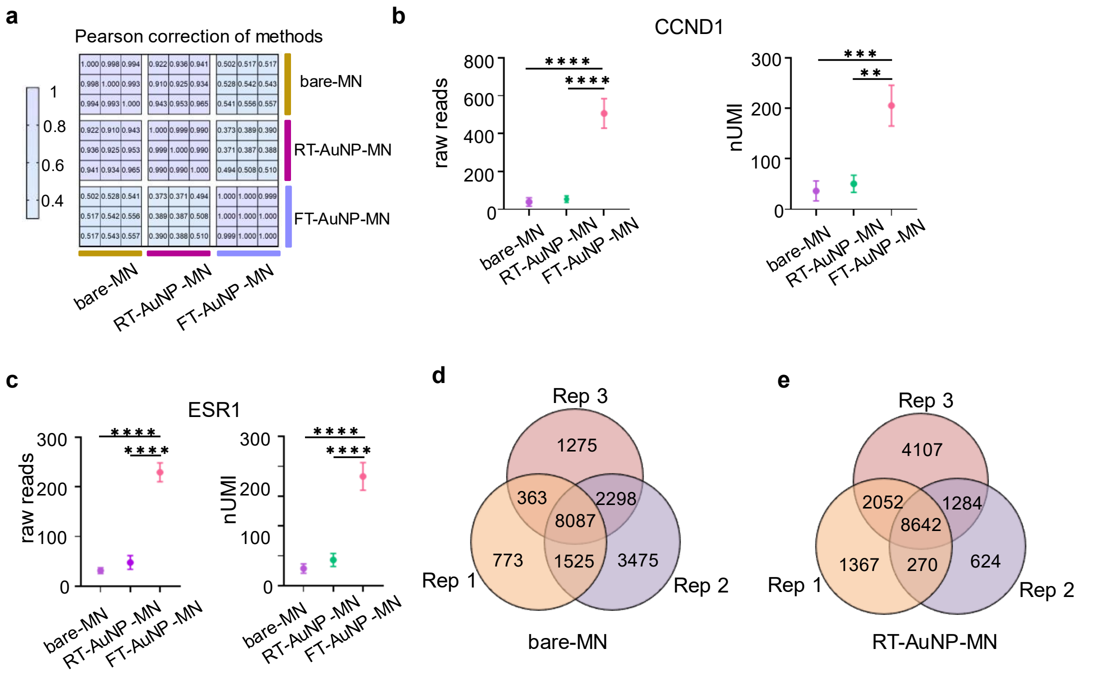

**Supplementary Fig. 3. Analysis the bare-MN, RT-AuNP-MN and FT-AuNP-MN.** (a) Comparison of bare-MN, RT-AuNP-MN and FT-AuNP-MN method using Pearson correlation analysis. (b) The raw reads count and nUMI of *CCND1* and (c) *ESR1*gene detected in MCF-7 cell ($n$ = 3 in each group). Data are presented as mean ± SD. (d) Genes detected in three replicates of bare-MN and (e) RT-AuNP-MN. Numbers of detected Genes are listed.

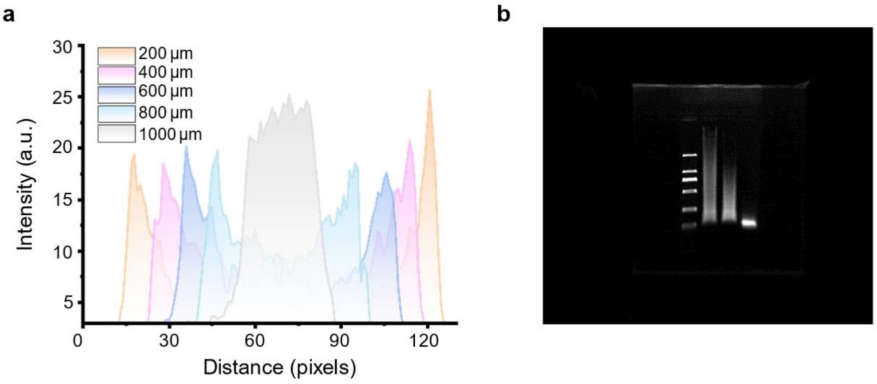

**Supplementary Fig 4. Characterization of fluorescently labeled single MN.** (a) Fluorescence intensity distribution profiles across different layers. (b) Full-size gel for Fig. 4h.

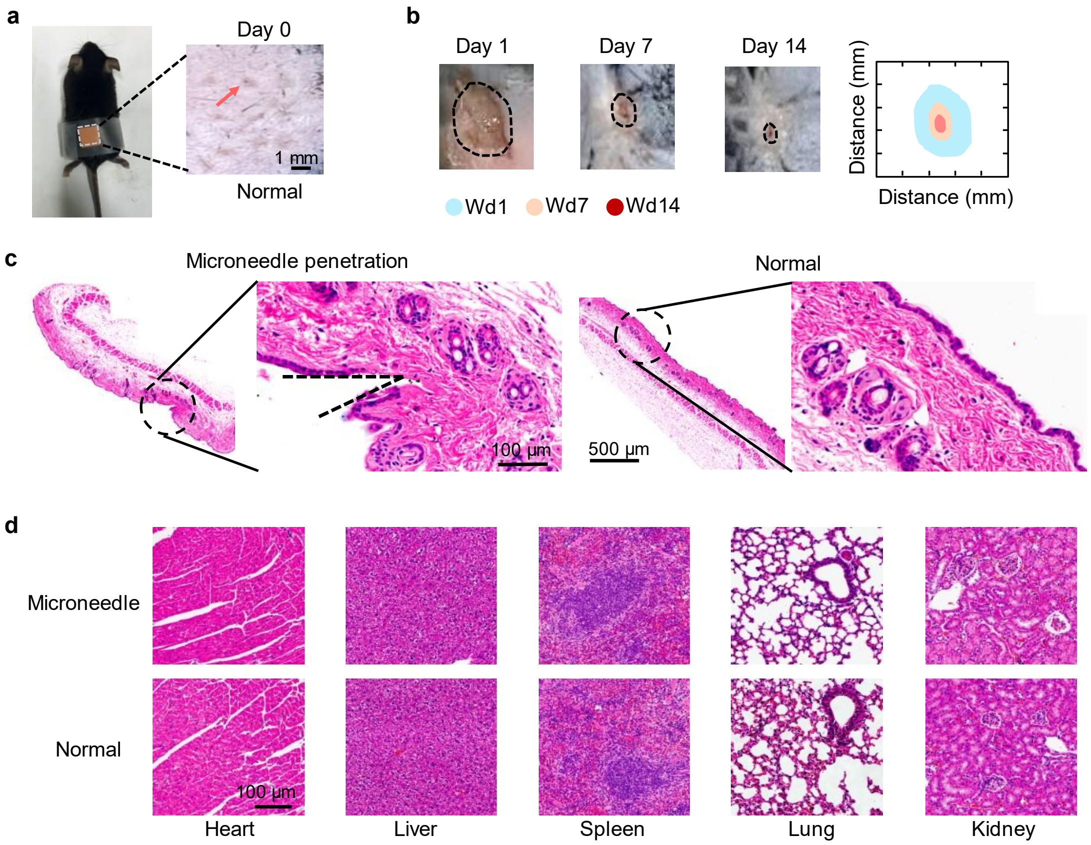

**Supplementary Fig. 5. Biophysical properties of MN patch.** (a) Mouse dorsum skin (the area within the white dashed line) was sampled using a MN patch, and the inset shows a magnified view of the MN patch penetrating the skin (The red arrow indicates the MN hole). Scale bar, 1 mm. (b) Optical photos and simulated graphs of wound closure changes at different time points (day 0, 1, 7 and 14). (c) H&E stained section of mouse skin showing the penetration of a single MN (left) and from the normal control group (right). Scale bar, 100 μm. (d) H&E stained section of mouse organs with and without administration of MN patch, including heart, liver, spleen, lung, kidney. Scale bar, 100 μm.

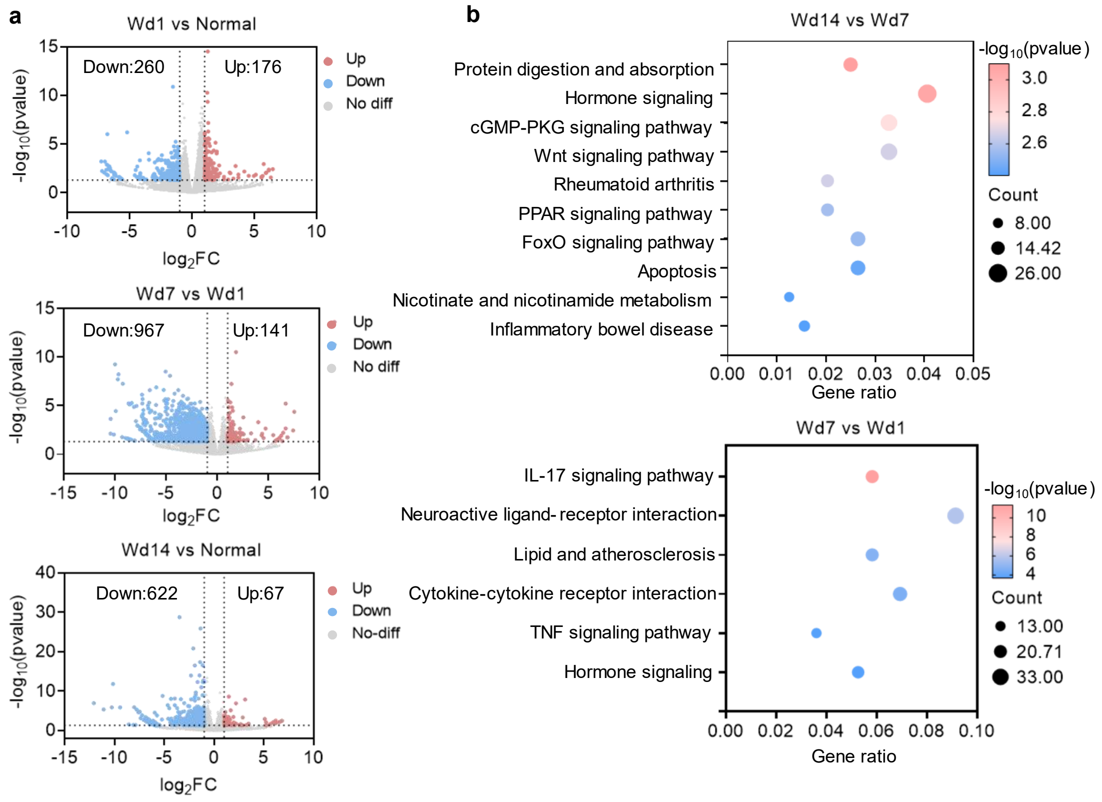

**Supplementary Fig. 6. Differential expression analysis of mouse skin wound healing by aivST-seq.** (a) Volcano plot showing differentially expressed genes between Wd1 vs Normal, Wd7 vs Wd1 and Wd14 vs Normal ($n$ = 3 in each group) (b) KEGG pathway enrichment analysis of Wd14 vs Wd7 and Wd7 vs Wd1 ($n$ = 3 in each group). Data are presented as mean ± SD.

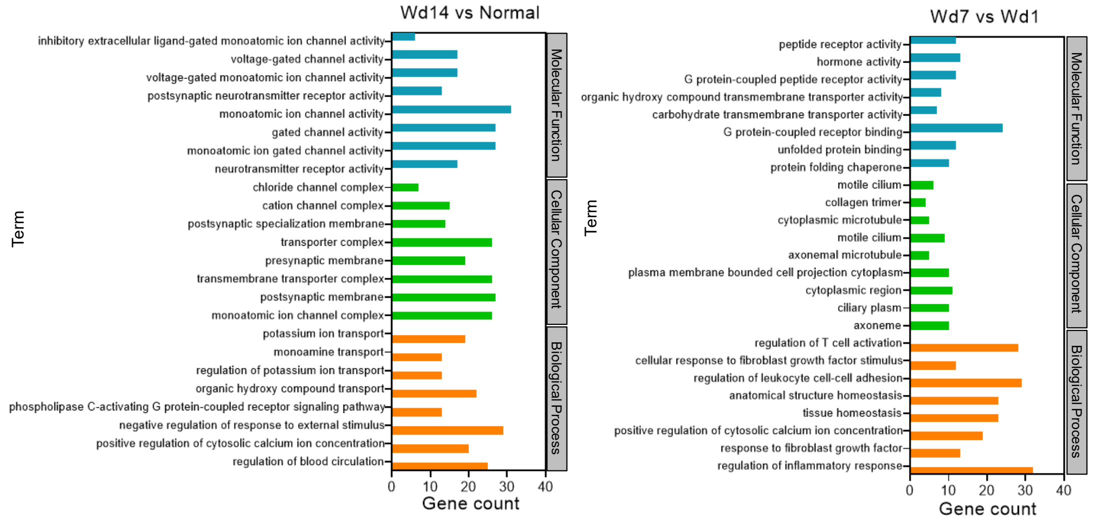

**Supplementary Fig. 7. GO analysis in Wd14 vs Normal and Wd7 vs Wd1.** Bar chart illustrating significant upregulated GO categories (including molecular function, cellular component and biological process) in Wd14 vs Normal and Wd7 vs Wd1 ($n$ = 3 in each group).

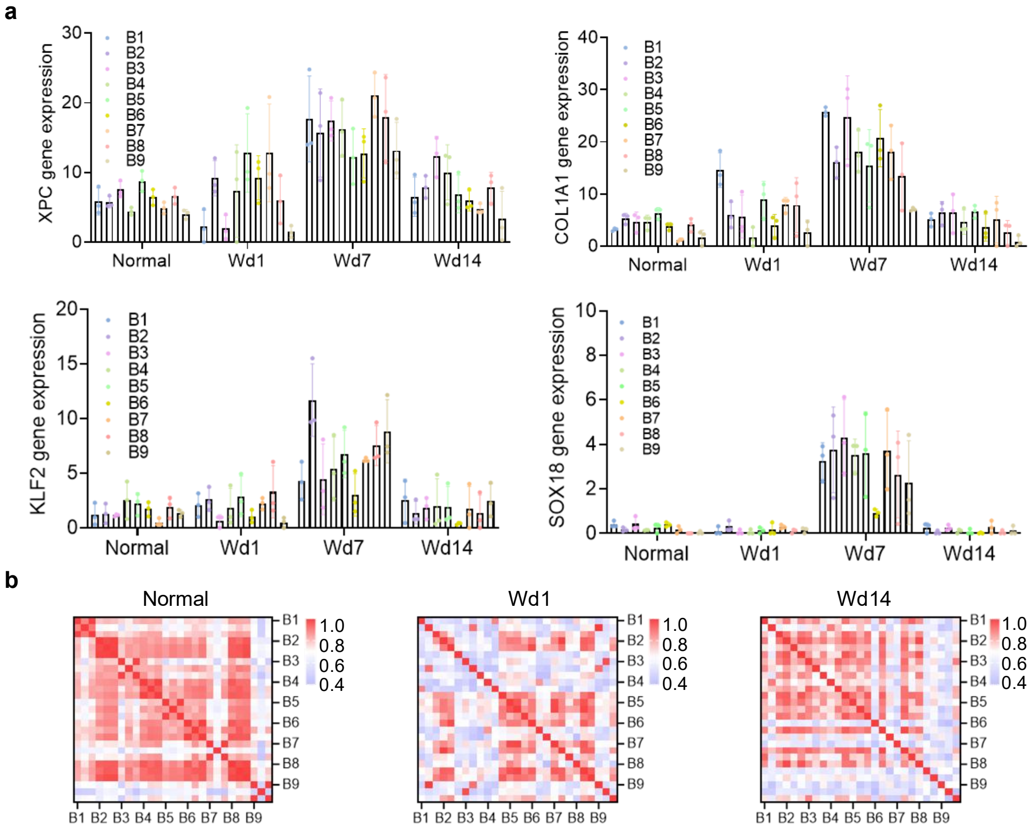

**Supplementary Fig. 8. Spatiotemporal dissection of wound healing using aivST-seq.** (a) Bar plot showing *XPC, COL1A1, KLF2* and *SOX18* gene normalized expression level at different spatial locations and at different conditions ($n = 3$ in each group). Data are presented as mean ± SD. (b) Heatmap showing pearson correction analysis of different barcodes in Normal, Wd1 and Wd14.
